# Genetic Determinants of Telomere Length in African American Youth

**DOI:** 10.1101/310417

**Authors:** Andrew M. Zeiger, Marquitta J. White, Sam S. Oh, Jonathan Witonsky, Maria G. Contreras, Pagé C. Goddard, Eunice Y. Lee, Kevin L. Keys, Lesly-Anne Samedy, Jennifer R. Liberto, Angel C.Y. Mak, Joaquín Magaña, Oona Risse-Adams, Celeste Eng, Donglei Hu, Scott Huntsman, Sandra Salazar, Adam Davis, Kelley Meade, Emerita Brigino-Buenaventura, Michael A. LeNoir, Harold J. Farber, Kirsten Bibbins-Domingo, Luisa N. Borrell, Esteban G. Burchard

## Abstract

Telomere length (TL) is associated with numerous disease states and is affected by genetic and environmental factors. However, TL has been mostly studied in adult populations of European or Asian ancestry. These studies have identified 34 TL-associated genetic variants recently used as genetic proxies for TL. The generalizability of these associations to pediatric populations and racially diverse populations, specifically of African ancestry, remains unclear. Furthermore, six novel variants associated with TL in a population of European children have been identified but not validated. We measured TL from whole blood samples of 492 healthy African American youth (children and adolescents between 8 and 20 years old) and performed the first genome-wide association study of TL in this population. We were unable to replicate neither the 34 reported genetic associations found in adults nor the six genetic associations found in European children. However, we discovered a novel genome-wide significant association between TL and rs1483898 on chromosome 14. Our results underscore the importance of examining these genetic associations with TL in diverse pediatric populations such as African Americans.

## INTRODUCTION

Telomeres are DNA-protein structures composed of tandem hexamer repeat sequences (TTAGGG_n_) that cap the ends of each chromosome^1^. Telomeres play a vital role in maintaining DNA stability and integrity, and are therefore, critical for preserving genomic information^2,3^. With each mitotic division, a portion of telomeric DNA is lost. The cell enters senescence upon reaching a critical telomere length (TL) threshold^4^. TL has thus become an important biomarker of aging and overall health^5–8^. A complex interaction between genetic^9^ and non-genetic factors^10^ affects TL. While heritability estimates of TL range from 36% to 82%^11^, much is still unknown about genetic factors leading to variation in TL^12,13^.

Although epidemiological research in pediatric populations has linked TL to early life adversity^14^ and environmental exposures^15,16^, few studies have focused on the genetic determinants of TL in pediatric populations. In contrast, several genetic studies of TL in European and Asian adults have identified and replicated 34 genetic variants associated with TL^17–26^. Over 30 studies have used these variants as genetic proxies for TL through Mendelian randomization approaches to address reverse causation when examining association between TL and disease in diseased patients^17,27,28^. However, recent studies in Chinese newborns and European children have failed to replicate these variants, suggesting that they are not generalizable across age groups^29,30^. One study, by Stathopoulou *et al*., reported six novel genetic variants associated with TL in European children (age 4-18 years) not previously discovered in adult telomere studies^30^. Replication of these six genetic variants has not yet been attempted. Given that adult TL appears determined prior to adulthood^31^, further research in diverse pediatric populations is necessary to validate the existence of genetic effects on TL early in life.

Previous genetic studies of TL have been done almost exclusively in populations of European ancestry^32^, yet there is evidence that TL varies by race/ethnicity^32–34^. African Americans have been shown to have longer telomeres throughout life^34–36^ and a greater rate of telomere attrition than populations of European ancestry^37^. Population-specific differences in genetic variants have previously been shown across the genome^38^. Thus, it is possible that population-specific variation of genetic factors contributing to TL influences the difference in TL observed between populations of African and European ancestries^33^.

To further understand the relationship between genetic variants and TL, we performed the first large-scale genetic study of TL in African American children and adolescents (n=492) from the Study of African Americans, Asthma, Genes and Environments (SAGE). Herein, we analyze genome-wide genetic data to attempt validation of previously reported genetic associations with TL and identify genetic variants influencing TL in African American children and adolescents.

## RESULTS

### Study Population

Demographic information for the study population (n=492) is presented in Table 1. The age of participants ranged from 8 to 20 with a median age of 15.8 (IQR = 12.4, 18.3; Table 1). Median African ancestry was 0.81 (IQR = 0.74, 0.85; Table 1) and increased African ancestry was significantly associated with longer TL (β = 0.333, P = 0.022, Figure 1). While individuals with public health insurance had significantly longer TL than individuals with private health insurance (P = 1.84 × 10^−4^), there was no significant association of age or maternal education with TL (Supplementary Table S1).

**Figure 1:**
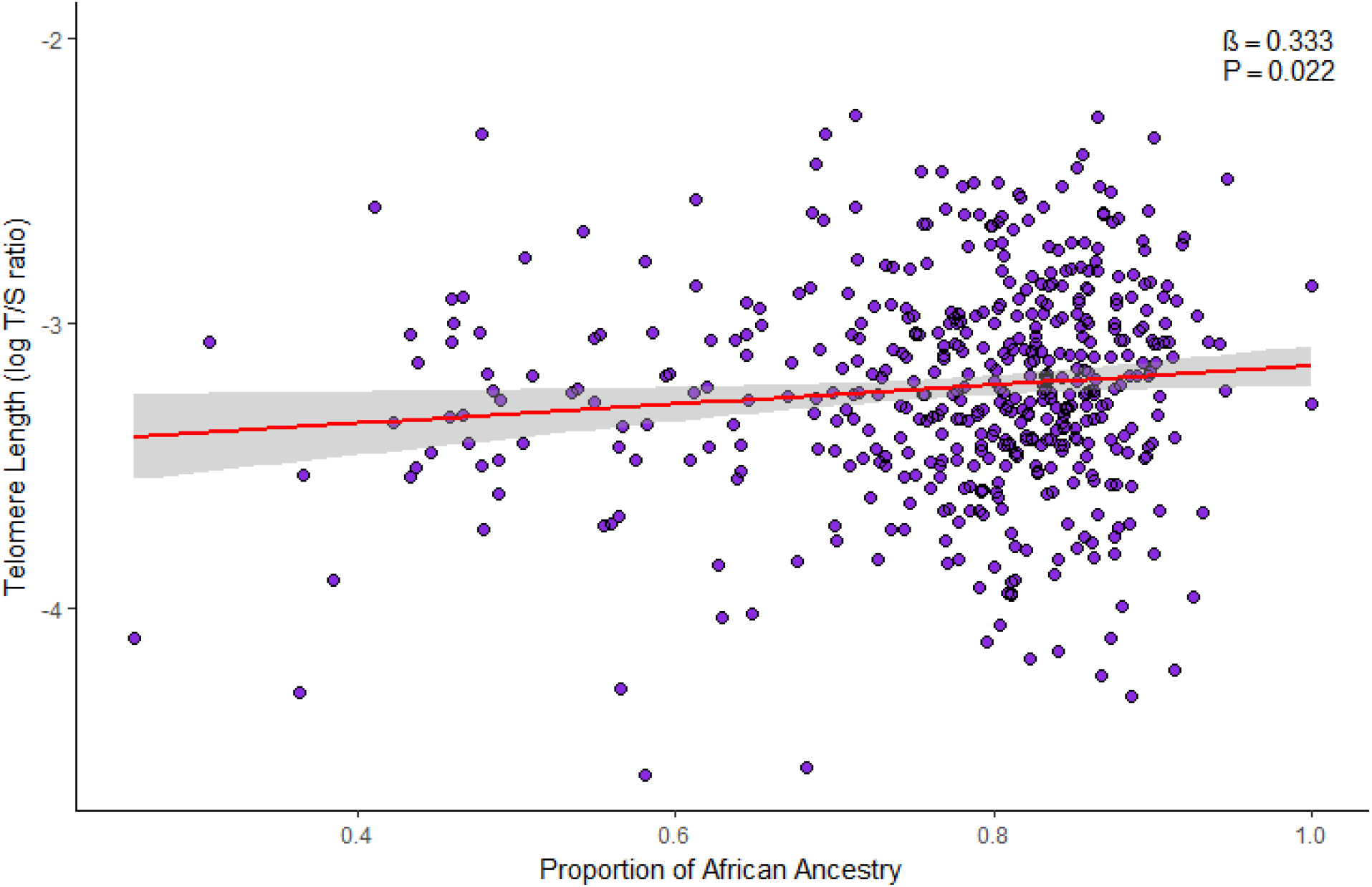
Association between log-transformed TL and African ancestry in healthy African American children and adolescents in SAGE: San Francisco Bay Area, 2006–2015.

**Table 1.**
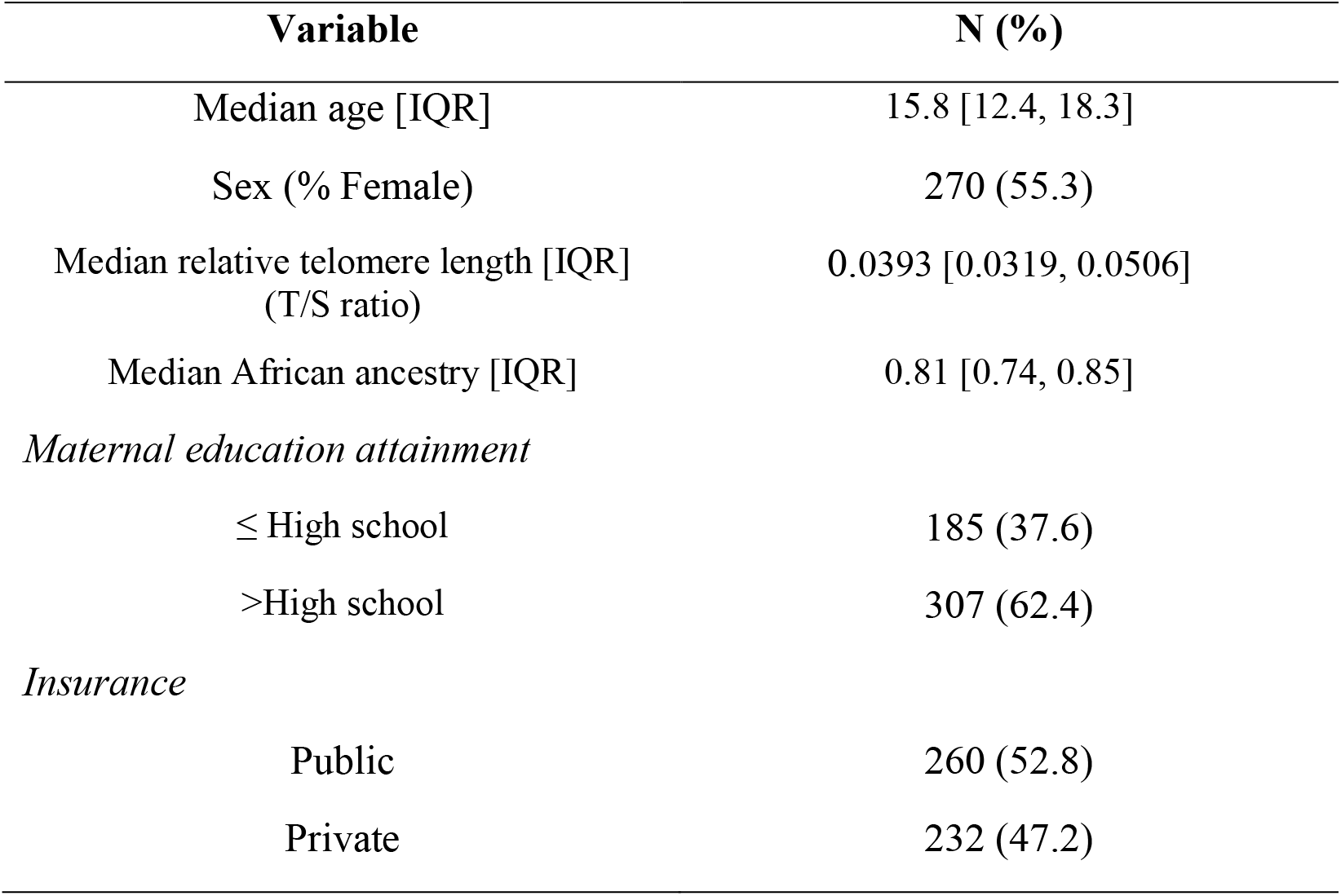
Demographic characteristics of healthy African American children and adolescents (n=492) in SAGE: San Francisco Bay Area, 2006–2015.

### Evaluation of Previous Variants

We evaluated 40 variants, 34 from adult studies (Table 2) and six from a pediatric study (Table 3), for an association with log-transformed TL. None of the variants from either the adult or pediatric studies were significantly associated with TL in our study population (P > 0.05). To determine whether the combined effect of the six previously discovered pediatric variants was associated with TL in our study population, we calculated a weighted genetic prediction score (GPS) by aggregating the allele associated with longer TL in European children weighted by the published β-coefficient^30^. There was no significant association between the GPS and TL in our study population of African American children and adolescents (β = 0.377, P = 0.150, Figure 2).

**Figure 2:**
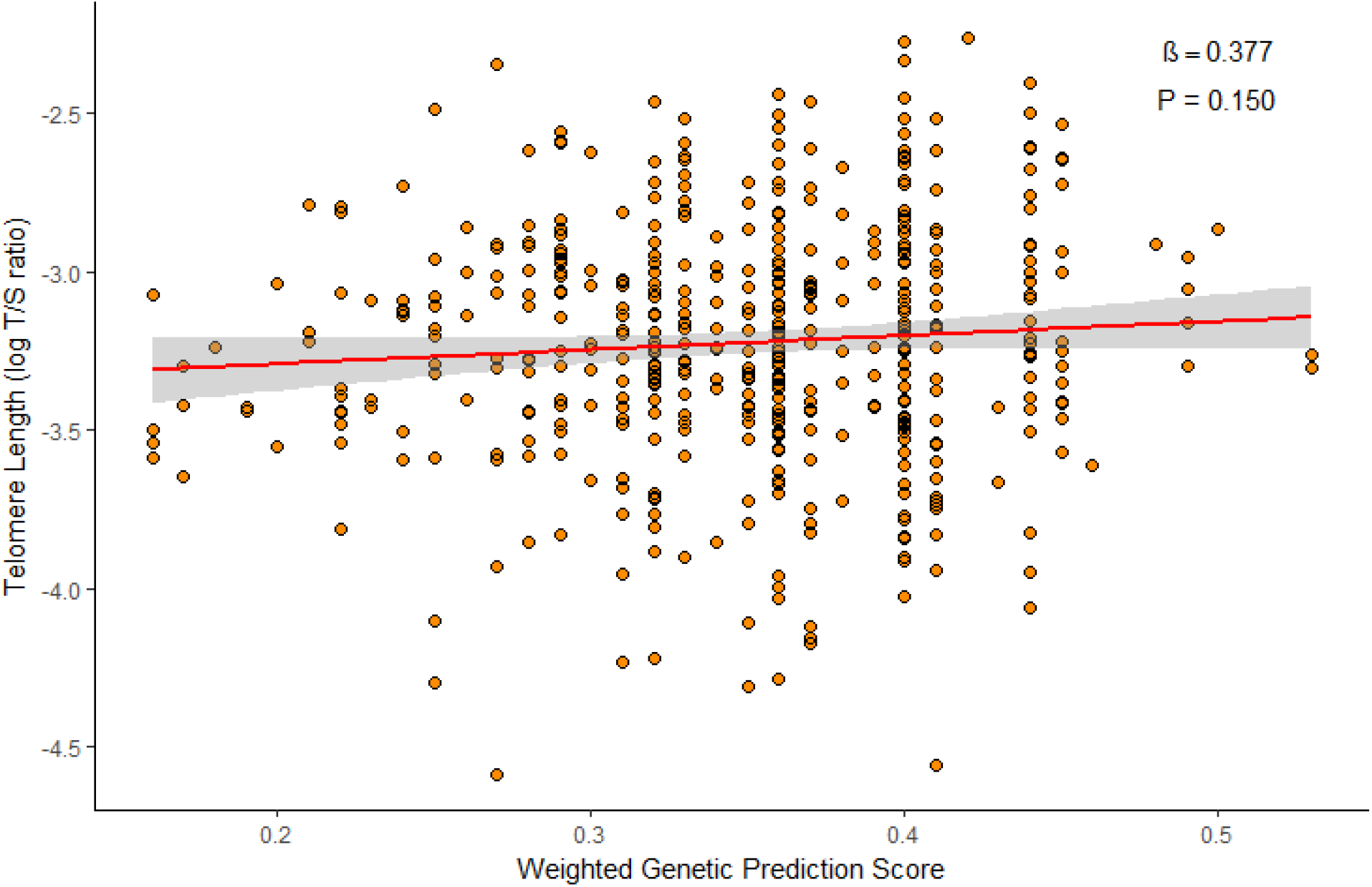
Adjusted association between log-transformed TL and GPS in healthy African American children and adolescents in SAGE: San Francisco Bay Area, 2006–2015. Regression association adjusted for sex, age, genetic ancestry, maternal educational attainment, health insurance type and batch effects.

**Table 2.**
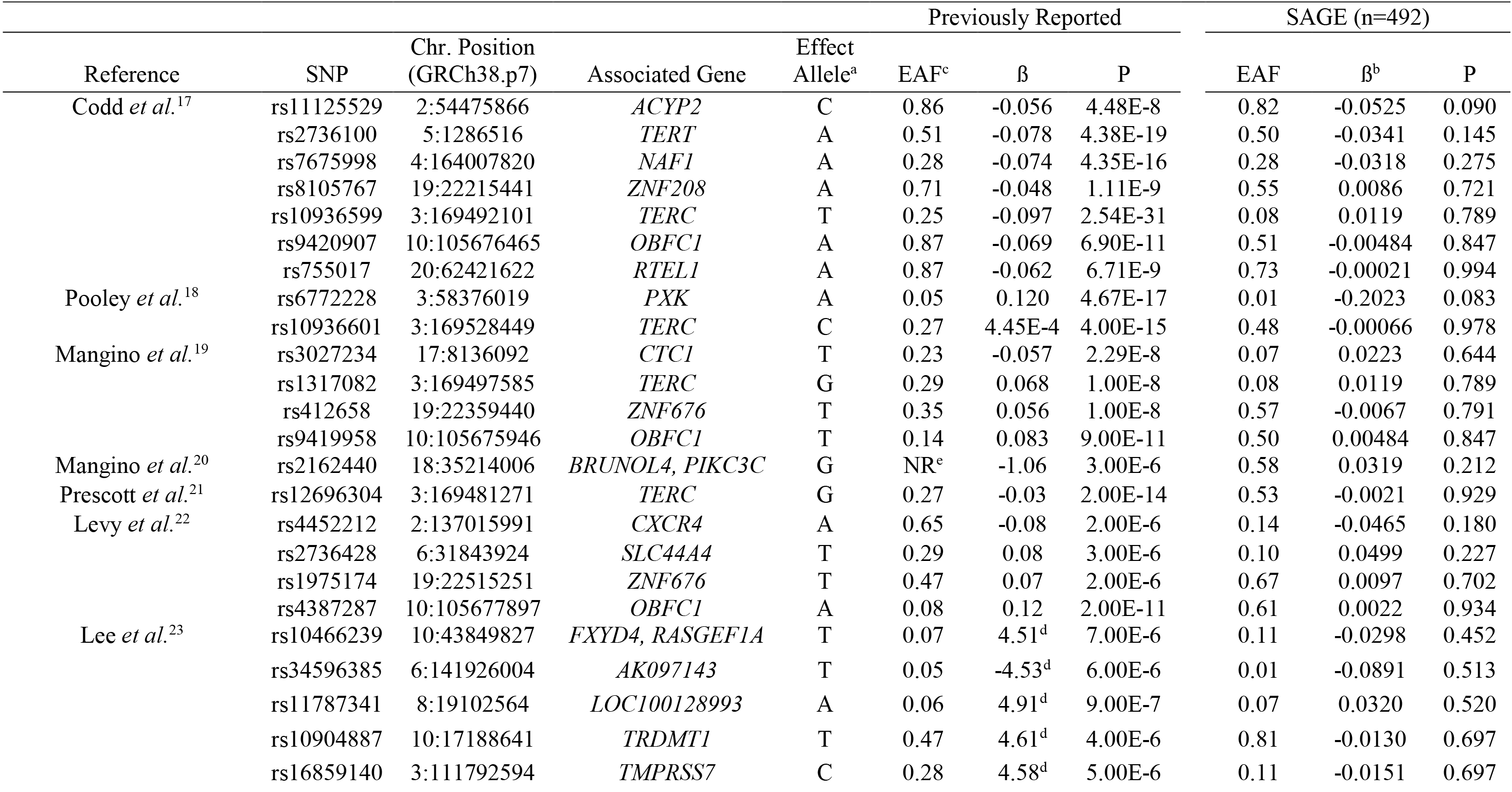
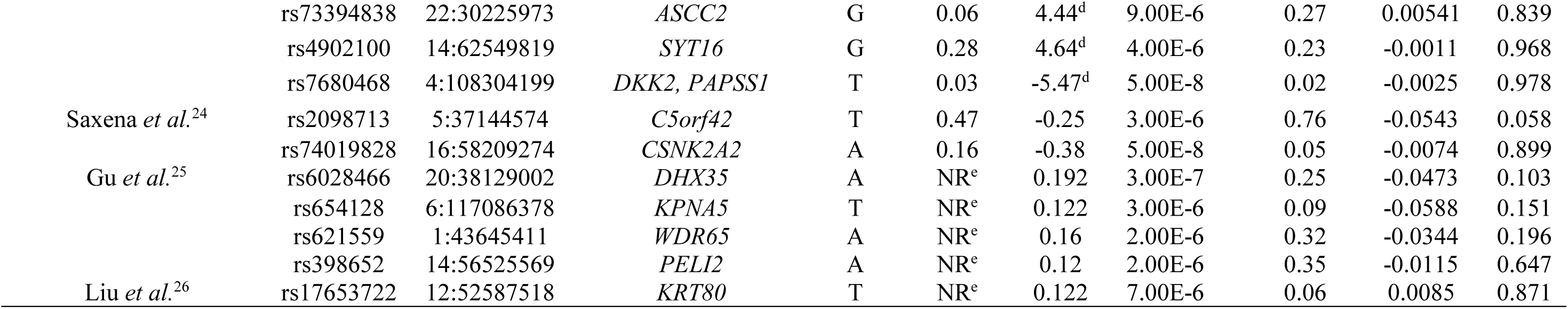
Adjusted analysis of log-transformed TL using 34 SNPs found in adult studies in healthy African American children and adolescents in SAGE: San Francisco Bay Area, 2006–2015. Regression adjusted for sex, age, genetic ancestry, maternal educational attainment, health insurance type and batch effects. ^a^Reported effect allele. ^b^Additive Linear regression β coefficient. ^c^EAF: Effect Allele Frequency in previous studies and current study. ^d^T-test test statistic reported instead of β coefficient. ^e^NR = Not Reported.

**Table 3.** Adjusted analysis of log-transformed TL using six SNP’s found in pediatric study healthy African American children and adolescents in SAGE: San Francisco Bay Area, 2006–2015. Regression adjusted for sex, age, genetic ancestry, maternal educational attainment, health insurance type and batch effects. ^a^Effect Allele. ^b^Additive Linear Regression β coefficient. ^c^EAF: Effect Allele Frequency. ^d^T-Test value used instead of β coefficient.

| SNP | Chr. Position | Associated Gene | EA<sup>a</sup> | Previously Reported |  |  | SAGE (n=492) |  |  |
| --- | --- | --- | --- | --- | --- | --- | --- | --- | --- |
| | | | | EAF<sup>c</sup> | $\beta$ (SE) | P | EAF<sup>c</sup> | $\beta^b$ (SE) | P |
| rs594119 | 6:124940213 | *NKAIN2* | T | 0.19 | -0.05 (0.01) | 2.19E-5 | 0.15 | -6.1E-2 (0.03) | 0.055 |
| rs11703393 | 22:44512124 | *PARVB* | G | 0.27 | -0.04 (0.01) | 1.69E-4 | 0.43 | -3.4E-2 (0.02) | 0.166 |
| rs2300383 | 21:35250086 | *ITSN1* | G | 0.47 | -0.04 (0.01) | 7.42E-6 | 0.73 | 2.2E-2 (0.03) | 0.438 |
| rs528983 | 4:115976690 | *NDST4* | G | 0.11 | -0.07 (0.01) | 7.88E-6 | 0.15 | -2.1E-2 (0.03) | 0.523 |
| rs12678295 | 8:2748377 | *MYOM2, CSMD1* | G | 0.40 | -0.04 (0.01) | 1.92E-4 | 0.22 | -4.7E-3 (0.03) | 0.870 |
| rs10496920 | 2:142996517 | *LRP1B, LOC100129955* | G | 0.18 | 0.05 (0.01) | 4.60E-4 | 0.15 | -4.4E-3 (0.03) | 0.895 |

### Discovery Genome-Wide Association Study

We performed a discovery GWAS to identify significant and suggestive associations between common genetic variants and TL in our study population. We identified a novel association between rs1483898 and TL that reached genome-wide significance (P = 7.86 × 10^−8^, Figure 3). Rs1483898 is an intergenic single nucleotide polymorphism (SNP) located proximal to the *LRFN5* gene on chromosome 14. An increase in copies of the rs1483898 A allele was significantly associated with longer TL (β = 0.148, P = 7.86 × 10^−8^, Figure 4). We also discovered 41 suggestive associations between common variants and TL (P < 2.32 × 10^−6^, Supplementary Table S2). Of particular note were rs9675924 (β = −0.171, P = 2.27 × 10^−6^, Supplementary Table S2) located in *CABLES1* and rs4305653 (β = 0.167, P = 1.81 × 10^−6^, Supplementary Table S2) located in *TTC37*. These genes have been previously associated with telomere biology^39,40^.

**Figure 3:**
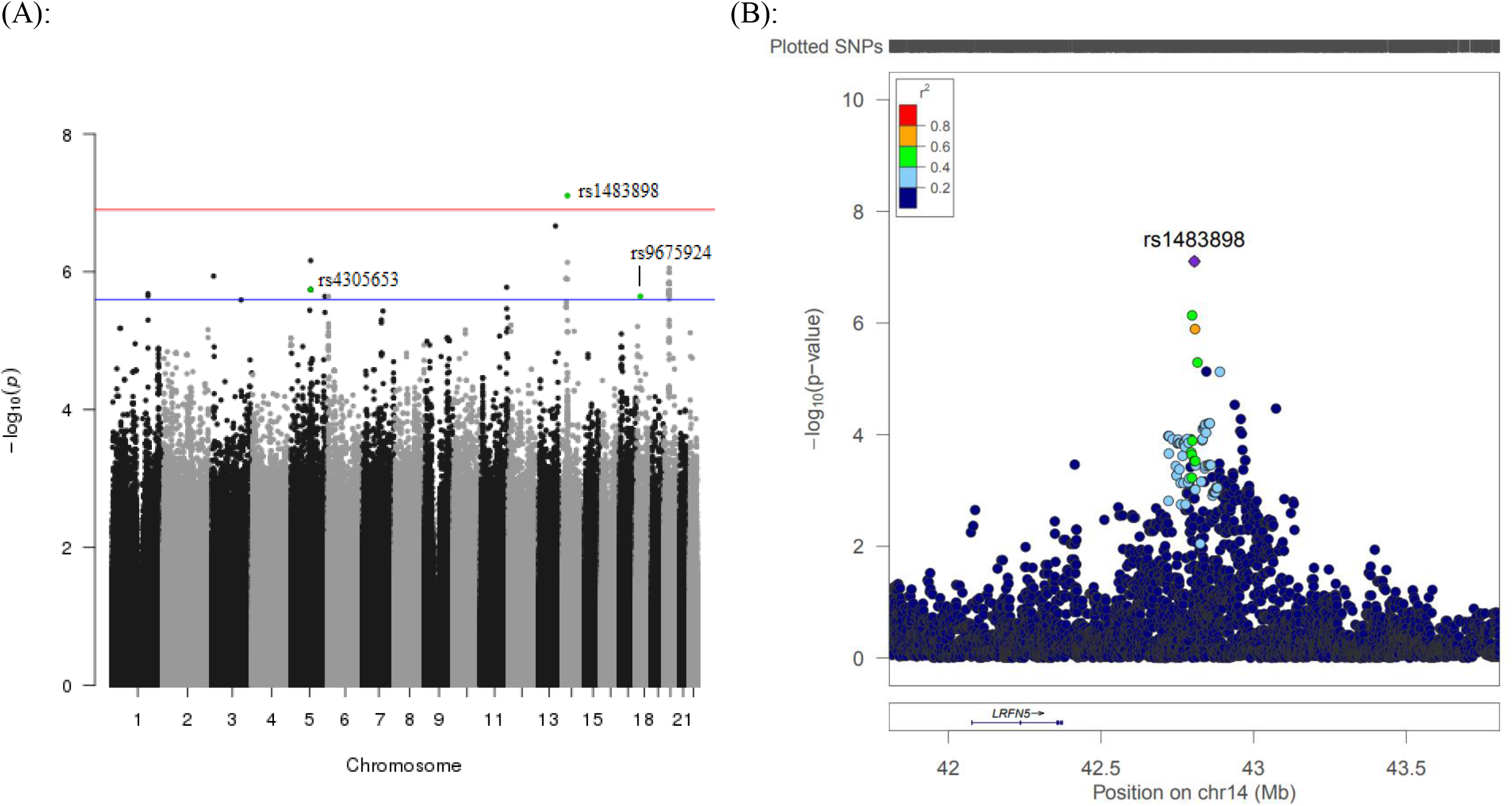
Results of GWAS for TL in healthy African American children and adolescents in SAGE: San Francisco Bay Area, 2006–2015. (A) Manhattan plot of the GWAS of TL with three SNPs relevant to telomere biology highlighted. Genome-wide significance threshold is indicated as red line (P = 1.2 × 10^−7^) and suggestive significance threshold is indicated as blue line (P = 2.3 × 10^−6^). (B) Expansion of 1 Mb flanking region around the top hit (rs1483898) with surrounding SNPs colored by amount of linkage disequilibrium with the top SNP, indicated by pairwise r^2^ values from hg19/November 2014 1000 Genomes AFR.

**Figure 4:**
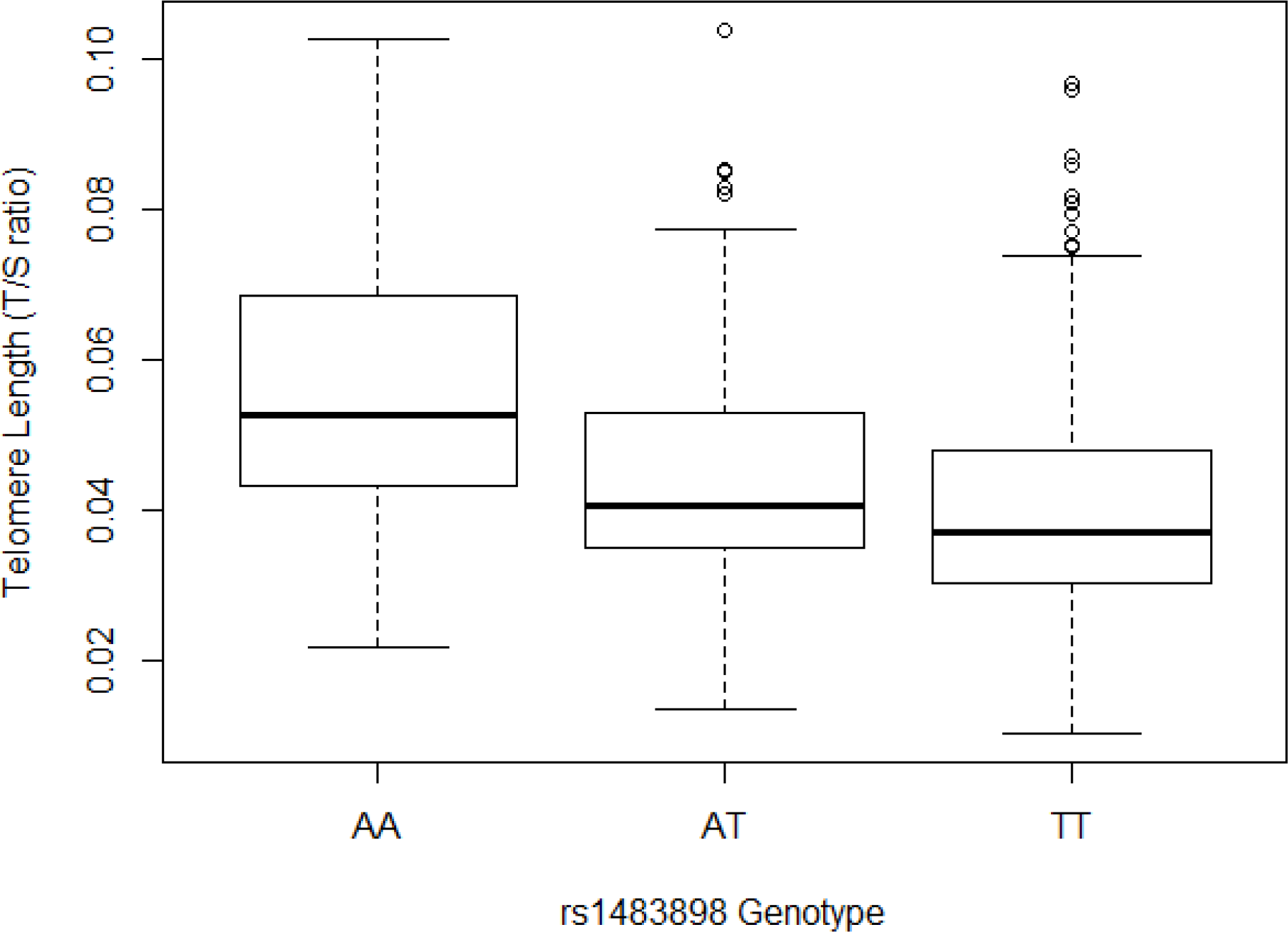
Comparison of mean TL between rs1483898 genotypes in healthy African American children and adolescents in SAGE: San Francisco Bay Area, 2006–2015.

## DISCUSSION

In this study, we contribute to the nascent body of research on genetic determinants of TL by assessing the generalizability of genetic markers of TL to African American children and adolescents. Our results are consistent with recent studies in pediatric populations^29,30^ that did not replicate variants associated with TL in adults^17–26^, suggesting that these variants may not play a significant role in the regulation of TL during the first two decades of growth and development. However, we were also unable to replicate genetic variants associated with TL in a population of European children^30^, highlighting potential population-specific effects of genetic associations with TL. Lastly, we identified a genome-wide significant variant, rs1483898, and 41 suggestively associated variants within genes relevant to telomere biology in a GWAS for TL in African American children and adolescents.

Genetic determinants of TL are critical to understanding inter-individual variation in TL. However, most studies of TL have been performed in adults, after the developmental time window when age-dependent telomere shortening may have already occurred^32^. Studies conducted among adults have identified and replicated 34 variants that have been used in recent years as proxies of TL in studies of disease risk^17,27^. We did not replicate these genetic associations in our study population of African American children and adolescents. Pediatric studies of TL by other groups^29,30^ have also been unable to replicate associations found among adults, suggesting that the genetic components influencing TL may differ between adult and pediatric populations. It is possible that variants identified in adults relate to telomere maintenance in adulthood but do not regulate TL during earlier developmental windows. For example, resistance to telomere shortening during childhood may be influenced by genetic factors impacting telomerase, a critical enzyme in telomere elongation^41^ that is influenced by genetic loci^42^ and shows age-related reduction in activity^43^. TL is determined prior to adulthood dependent on the TL setting at birth and the rates of shortening and elongating during the first two decades of life^10^. These factors have genetic influences that have yet to be fully characterized^1,10^.

We attempted replication of six TL-associated genetic variants discovered in healthy children of European ancestry^30^. We found no significant association with TL among the six variants independently or in a weighted GPS, which tests cumulative variation at multiple genetic loci. Heritability estimates of TL range from 36% to 82%^11^, yet has only been reported in populations of European ancestry and may not be generalizable to other populations. Similarly, genetic determinants of TL have primarily been studied in populations of European or Asian descent. Recent studies attempting to replicate and/or identify genetic associations with TL in non-European populations, including Punjabi Sikh^24^, Han Chinese^26,44^ and Bangladeshi^45^, have had mixed success. Among the limited set of studies assessing TL-associated genetic variants in populations of African ancestry, all have been performed in adult populations. One study discovered genetic variants associated with TL in adults of European ancestry that were not associated in an adult population of African Americans^22^. Another study in adult African Americans was only able to replicate the effects of variants in *TERC*, the gene encoding the enzyme telomerase, that had been identified in populations of European ancestry^46^. We found a significant positive association between the proportion of African ancestry and TL in African American children and adolescents, which is consistent with research among adults^34^. Considering TL dynamics vary by race/ethnicity^32–34^, our study augments the current literature by demonstrating that TL-associated genetic variants differ between ancestral populations in the pediatric age range. Ancestry-specific genetic associations with a phenotypic trait have been demonstrated previously^47^, thus, the difference we observed may result from population-specific effects impacting genetic regulation of TL. It is worth noting regional variation in environmental and social exposures between SAGE’s urban San Francisco Bay Area and the more rural Nancy, France of the Stathopoulou *et al*. study^48^ as potential factors effecting the association between genetic variants and TL and possibly precluding replication of the Stathopoulou *et al*. study.

We identified associations with TL in biologically relevant pathways relating to apoptosis, cell senescence and telomere replication. The most significant association, reaching genome-wide significance, was for rs1483898. Rs1483898 is located on the q21.1 arm of chromosome 14 with the closest gene, *LRFN5*, encoding a neuronal transmembrane protein. In our population of African American children and adolescents, increasing copies of the A allele associated with longer TL. The A allele of rs1483898 has allele frequencies of 0.74, 0.86 and 0.45 in the African, European and East Asian populations of 1000Genomes, respectively^49^.

We identified 41 variants that were suggestively associated with TL. The A allele of rs9675924, located within cell cycle regulator *CABLES1* on chromosome 18, associated with shorter TL. *CABLES1* is co-expressed with protein kinase *CDK5*, a known contributor to apoptosis in certain neuronal diseases^39^. *CABLES1* has also been shown to inhibit cell proliferation and induce cell senescence in umbilical endothelial cells^40^. The C allele of rs4305653 associated with longer telomeres; the variant is located on chromosome 5 within *TTC37*, a component of the SKI complex which mediates protein-protein interactions. *TTC37* is co-expressed with apoptosis-promoting protein *APAF1* and with *TEP1*, a protein that binds to *TERC* and is essential for telomere replication^39^. Ultimately, co-expression is only a proxy for co-regulation^50^; replication and further investigation of our results are needed to better characterize relevant associations between these genetic loci and telomere biology.

Our study lacked an independent replication cohort to assess the reproducibility of our genetic associations due to the unique characteristics of our study population (African American children and adolescents with genetic and TL data). Measurement of TL in our study population provided a snapshot of TL at a specific point in the life course. Longitudinal studies of TL are required to understand changes in TL over the life course. Our inability to replicate the reported variants may also be explained by limited statistical power to discover weak or moderate genetic effects on TL. The major advantages of our study are that (1) it is the first large-scale study to investigate the genetic determinants of TL in a population of minority children and adolescents, and (2) our depth of phenotype data allowed us to adjust for social, environmental and genetic covariates (Supplementary Table S1).

In summary, the paucity of research on factors affecting TL in pediatric and non-European populations creates a knowledge gap in the scientific understanding of gene-environment interactions regulating telomeres. Epidemiological studies reporting associations between TL and disease risk are potentially biased by the disease itself or exposures relating to treatment. Genetic proxies for TL have recently been employed to overcome these and other potential biases, such as social and environmental exposures. A critical assumption when using genetic proxies for TL is that they are generalizable across age and racial/ethnic groups. However, we were unable to replicate previous findings of TL-associated variants in our study population. We also identified novel genetic associations with TL that have not been identified in previous studies in pediatric or adult populations. Further telomere research in pediatric populations from diverse ancestral backgrounds is required to fully understand the impacts of age- and population-effects on the genetic regulation of TL.

## METHODS

### Ethics statement

This study has been approved by the institutional review boards of University of California San Francisco and all participant centers. Written informed consent was obtained from all subjects or from their appropriate surrogates for participants under 18 years old.

### Study population

Our study included 492 healthy controls from the Study of African Americans, Asthma, Genes and Environments (SAGE). SAGE is one of the largest ongoing gene-environment interaction studies of asthma in African American children and adolescents in the USA. SAGE includes detailed clinical, social, and environmental data on asthma and asthma-related conditions. Full details of the SAGE study protocols have been described in detail elsewhere^51–53^. Briefly, SAGE was initiated in 2006 and recruited participants with and without asthma through a combination of clinic- and community-based recruitment centers in the San Francisco Bay Area. Recruitment for SAGE ended in 2015. All participants in SAGE self-identified as African American and self-reported that all four grandparents were African American.

After all quality control procedures relating to TL measurement, TL was computed for 596 healthy controls in SAGE from whole blood. There were 495 healthy controls with complete sex, age, African ancestry, maternal educational attainment and health insurance information. Three individuals showed extreme outlier measurements for TL (three times the interquartile range) and were thus removed.

### Covariates

Maternal educational attainment and health insurance type were used as proxies of SES^54–56^. Maternal educational attainment was dichotomized based on whether a participant’s mother had pursued education beyond high school (i.e., ≤12 versus >12 years of education). Health insurance type was defined as private versus public insurance. The genetic ancestry of each participant was determined using the ADMIXTURE software package^57^ with the supervised learning mode assuming two ancestral populations (African and European) using HapMap Phase III data from the YRI and CEU populations as references^58^.

### Variant selection and genotyping

TL associated variants were selected for replication using the following criteria set *a priori*: i) published association reaches genome-wide significance (P ≤ 5 × 10^−8^) on NHGRI-EBI GWAS Catalog by October 26, 2017; ii) variant used as genetic proxy of TL in at least one study; iii) variant reaches suggestive genome-wide significance (P ≤ 5 × 10^−5^) in a novel GWAS of TL in children; iv) variant has a minor allele frequency (MAF) of at least 1% in the SAGE study population. We identified variants from 11 studies^17–26, 30^. Ten of the 11 studies were performed in adult populations and nine of the 11 studies were performed in populations of European descent, with the remaining two performed in Punjab Sikh^24^ and Han Chinese^26^ populations. In total, we identified 40 variants from the literature, of which 12 were genotyped and 28 were imputed. The 28 imputed SNPs had r^2^ (squared correlation between imputed and expected genotypes) ranging from 0.88 to 1.00.

DNA was isolated from whole blood collected from SAGE participants at the time of study enrollment using the Wizard^®^ Genomic DNA Purification kits (Promega, Fitchburg, WI). Samples were genotyped with the Affymetrix Axiom^®^ LAT1 array (World Array 4, Affymetrix, Santa Clara, CA), which covers 817,810 SNPs. This array was optimized to capture genetic variation in African-descent populations such as African Americans and Latinos^59^. Genotype call accuracy and Axiom array-specific quality control metrics were assessed and applied according to the protocol described in further detail in Online Resource 1. Data was submitted to the Michigan Imputation Server and phased using EAGLE v2.3 and imputed from the Haplotype Reference Consortium r1.1 reference panel using Minimac3^60^. Imputed SNPs were included if they had an r^2^ higher than 0.3. Quality control inclusion criteria consisted of individual genotyping efficiency > 95%, Hardy-Weinberg Equilibrium (HWE) P > 10^−4^, and MAF > 5%. Cryptic relatedness was also assessed to ensure that samples were effectively unrelated. Samples with an estimate of genetic relatedness greater than 0.025 were excluded. After quality control procedures, 7,519,176 imputed and genotyped SNPs were available for analysis.

### Telomere length measurement

#### DNA isolation and quantification

Genomic DNA was isolated from whole blood according to manufacturer’s recommendation using Wizard^®^ Genomic DNA Purification Kits (Promega, Fitchburg, WI). A NanoDrop^®^ ND-1000 spectrophotometer (Thermo Scientific) was used to assess DNA quality and quantity. All samples assayed had absorbance ratios (260/280) between 1.8 and 2.0.

#### Determination of Relative Telomere Length

Relative TL for each sample was determined using the quantitative real time PCR (qPCR) method first described by Cawthon *et al*., which quantified TL in terms of telomere/single copy gene (T/S) expression ratios^61^. This protocol was modified with regard to data processing and control samples as previously published by O’Callaghan *et al*. and described in further detail in Supplemental Methods^62^. In brief, relative TL for each sample was calculated using the delta-delta C_T_ (2^−ΔΔCt^) formula^61^. Using this formula, the TL computed for each SAGE sample is proportional to the T/S ratio of that sample normalized to the T/S ratio of the PCR plate positive DNA control sample^61,63,64^. Inter- and intra-experimental coefficients of variation for our internal control (1301 cell line DNA) were 3% and 4.25%, respectively. Average amplification efficiency across plates was ≥ 90% for telomere and 36B4 assays. As TL was not normally distributed in our study population, we performed all parametric tests on a log-transformation of TL.

### Replication analysis

Genotypes for all 40 previously published SNP’s in adults and children were tested for association with log-transformed TL in a multivariable linear regression analysis. Regression analyses were run separately for each SNP under an additive model to calculate the individual effect of the SNP on TL. Each regression analysis was adjusted for biological, environmental and social factors that may impact TL including sex, age, African ancestry, maternal education, and health insurance type. We adjusted for qPCR plate ID in all regression analyses to ensure that our results were not impacted by qPCR batch effects. To ensure direct comparison of results between previous studies and our current study we coded the effect alleles in our analysis to be the same as those used in previous studies.

### Genetic Prediction Score construction

Recent research suggests the cumulative effect of multiple genetic markers may be a stronger predictor of a quantitative phenotype than the individual markers^65,66^. We therefore constructed a weighted GPS based on the six variants from Stathopoulou *et al*. to test their cumulative effect on TL^30^. We calculated each subject’s weighted GPS by summing the number of alleles (0, 1 or 2) associated with longer telomeres after weighting the allele count by the reported β-coefficient from the literature. We assumed that an effect allele having a positive β-coefficient meant that each additional copy of that allele was positively associated with TL. We used the GPS as a predictor in a linear regression against log-transformed TL controlling for sex, age, genetic ancestry, maternal educational attainment, health insurance type and batch effects. We were unable to calculate a weighted GPS based on the 34 variants in adult studies because the effect size could not be standardized across the studies.

### Calculation of population-specific genome-wide significance threshold

The standard GWAS threshold for statistical significance is 5 × 10^−8^. This number was derived by applying a Bonferroni correction for multiple testing to a dataset of one million independent markers/SNPs. However, in many cases, this threshold is overly conservative and can be inappropriate when (1) a smaller number of markers is genotyped, and (2) the assumption of independence of tests is violated.

In order to adjust the Bonferroni correction based on the actual number of independent test performed on our dataset, we determined the number of independent tests using the protocol published by Sobota *et al*.^67^ This method estimates the effective number of independent tests in a genetic dataset after accounting for linkage disequilibrium (LD) between SNPs using the LD pruning function in the PLINK 1.9 software package^68^. The following parameters were used in PLINK 1.9 as advised by the authors: 100 SNP sliding window, step size of 5 base pairs, and a variance inflation factor of 1.25. Applying this method on 7,519,176 genotyped and imputed SNPs yielded 431,896 independent tests, which was then used to calculate the genome-wide significance threshold (Bonferroni correction 0.05/431,896=1.2 × 10^−7^). A suggestive threshold was set at 2.3 × 10^−6^ for association results based on the widely used formula: 1/(effective number of tests)^69^.

### Discovery Genome-Wide Association Study

We performed a genome-wide association study (GWAS) using 7,519,176 genotyped and imputed SNPs to assess the relationship between SNP genotype and log-transformed TL. The GWAS linear regression model adjusted for sex, age, African ancestry, maternal educational attainment, health insurance type and batch effects. All testing was performed using PLINK1.9^68^. Manhattan plots (Figure 3a, 3b) were generated using the qqman package^70^ in the R statistical software environment (R Development Core Team 2010) and LocusZoom^71^. Curated protein-protein interactions were extracted using the STRING database^39^. An integrated confidence score for the interaction ranges from 0.5 (medium confidence) to 1 (high confidence).

## ACKNOWLEDGEMENTS

The authors acknowledge the patients, families, recruiters, health care providers and community clinics for their participation. In particular, the authors thank clinical recruiter Lisa Caine, RT. This work was supported in part by the Sandler Family Foundation, the American Asthma Foundation, the RWJF Amos Medical Faculty Development Program, the Harry Wm. and Diana V. Hind Distinguished Professor in Pharmaceutical Sciences II, the National Heart, Lung, and Blood Institute (NHLBI) R01HL117004, R01HL128439, R01HL135156, X01HL134589, the National Institute of Environmental Health Sciences R01ES015794, R21ES24844, the National Institute on Minority Health and Health Disparities P60MD006902, R01MD010443, RL5GM118984 and the Tobacco-Related Disease Research Program under Award Number 24RT-0025. M.J.W. was supported by a diversity supplement of NHLBI R01HL117004, an Institutional Research and Academic Career Development Award K12GM081266, and a NHLBI Research Career Development (K) Award K01HL140218. K.L.K. was supported by a diversity supplement of NHLBI R01HL135156. Research reported in this article was funded by the National Institutes of Health Common Fund and Office of Scientific Workforce Diversity under three linked awards RL5GM118984, TL4GM118986, 1UL1GM118985 administered by the National Institute of General Medical Sciences.

## AUTHOR CONTRIBUTIONS

A.M.Z, M.J.W, J.W., S.S.O, and E.G.B. were involved in the conception and design of the study. A.M.Z., M.J.W, S.S.O., E.Y.L., J.W., P.C.G., J.R.L., A.C.Y.M., C.E., D.H., S.H., M.G.C., L.A.S., K.L.K., O.R.A., J.M., and E.G.B were involved in the analysis and interpretation of data. S.S., C.E., A.D., K.M., E.B.B., M.A.L., H.J.F., K.B.D., L.N.B., and E.G.B. planned and supervised the collection of data. C.E., O.R.A., M.G.C., and M.J.W. generated telomere data. All authors provided revisions and approval of the final manuscript.

